# Motor learning in reaching tasks leads to homogenization of task space error distribution

**DOI:** 10.1101/2021.09.01.458189

**Authors:** Puneet Singh, Oishee Ghosal, Aditya Murthy, Ashitava Ghosal

**Affiliations:** John A. Paulson School of Engineering and Applied Sciences, Harvard University, Cambridge, MA, USA; Centre for Neuroscience, Indian Institute of Science, Bengaluru, India; Department of Mechanical Engineering, Indian Institute of Science, Bengaluru, India

**Keywords:** Motor learning, Reaching tasks, Force-field perturbation, Error along directions, Redundancy, Homogenization of error distribution

## Abstract

A human arm, up to the wrist, is often modelled as a redundant 7 degree-of-freedom serial robot. Despite its inherent nonlinearity, we can perform point-to-point reaching tasks reasonably fast and with reasonable accuracy in the presence of external disturbances and noise. In this work, we take a closer look at the task space error during point-to-point reaching tasks and learning during an external force-field perturbation. From experiments and quantitative data, we confirm a directional dependence of the peak task space error with certain directions showing larger errors than others at the start of a force-field perturbation, and the larger errors are reduced with repeated trials implying learning. The analysis of the experimental data further shows that a) the distribution of the peak error is made more uniform across directions with trials and the error magnitude and distribution approaches the value when no perturbation is applied, b) the redundancy present in the human arm is used more in the direction of the larger error, and c) homogenization of the error distribution is not seen when the reaching task is performed with the non-dominant hand. The results support the hypothesis that not only magnitude of task space error, but the directional dependence is reduced during motor learning and the workspace is homogenized possibly to increase the control efficiency and accuracy in point-to-point reaching tasks. The results also imply that redundancy in the arm is used to homogenize the workspace, and additionally since the bio-mechanically similar dominant and non-dominant arms show different behaviours, the homogenizing is actively done in the central nervous system.

**Significance:** The human arm is capable of executing point-to-point reaching tasks reasonably accurately and quickly everywhere in its workspace. This is despite the inherent nonlinearities in the mechanics and the sensorimotor system. In this work, we show that motor learning enables homogenization of the task space error thus overcoming the nonlinearities and leading to simpler internal models and control of the arm movement. It is shown, across subjects, that the redundancy present in the arm is used to homogenize the task space. It is further shown, across subjects, that the homogenization is not an artifact of the biomechanics of the arm and is actively performed in the central nervous system since homogenization is not seen in the non-dominant hand.

## Introduction

Movement kinematics during reaching movements are reasonably homogenous despite the presence of inhomogeneous biomechanics. This suggests that internal models which mediate sensorimotor transformations could play a role in mitigating such inhomegenities Click or tap here to enter text. while planning and executing movements (Mussa-Ivaldi et al. 1985). One such signature of such inhomegenieties is the observeddirection dependence of error in reaching tasks was done by (Gordon et al. 1994). who showd that the variability in the direction errors was larger along the axis of movement of the forearm than in the perpendicular direction, since the former requires greater forces to overcome the higher inertial load of the relatively larger shoulder movement. The dependence of task space error is also well-known in the robotics community -- in a serial robot with rotary joints, the position and orientation of an end-effector (hand) can be related to the joint variables in terms of trigonometric functions. As a consequence, the map between the hand velocities/error and the joint rates is nonlinear and the Jacobian matrix for a serial robot is not constant (Salisbury and Craig 1982). One of the consequence is that at any location in the workspace of a serial robot, the velocity distribution is an ellipse for 2D motion (an ellipsoid for 3D motion) and at different points in the workspace, the shape and size of the velocity ellipse (ellipsoid) will vary (Ghosal and Roth 1987; Salisbury and Craig 1982). In the robotics community, the velocity at a point in the workspace is often a proxy for the task space error as higher the velocity the less is the positioning accuracy. Although not explicitly mentioned and investigated in (Singh et al. 2016), a careful look at the results clearly show that the task space error is not the same in all directions (Singh et al. 2016) In this work, we look at the direction dependence of task space error and learning along the directions of reaching tasks performed by the human arm to show how internal models learn to homogenise the output.To get a better understanding of the positioning error in arm movement and its dependence with direction, in this work we take a re-look at the task space error in reaching tasks when the hand is subjected to a force-field perturbation. It is well-known that the magnitude of the task space error decreases and the central nervous system learns to adapt to the external force-field. In this work we go beyond and ask questions such as is there a directional dependence of error in reaching tasks when the hand motion is subjected to a force-field perturbation? Does the task space error reduction go to the level of error when there is no external force-field and finally what is the nature of the error distribution in the beginning and at the end of learning? We also ask the question if the redundancy in the human arm is used differently in the different directions and if the observed learning is actively controlled by the central nervous system.

## Materials and Methods

### Subjects

Twenty-two subjects (aged 22 ± 6 years) participated in the study. The handedness of the subjects was tested by modified Edinburgh Handedness Index (Salmaso and Longoni 1985). None of the subjects had any neurological diseases or chronic medication issues. All the subjects were paid for participation and gave informed consents in accordance with the institutional ethics committee of the Indian Institute of Science, Bangalore.

### Experimental setup

All the recordings were done in a dark room with the subjects sitting on chair with backrest and their chins resting on a chin rest with head locked with head bars on both sides of their temple as shown schematically in figure 1A. They looked down on a semi-transparent mirror on which they saw the targets while they moved a robotic arm handle on a plane below the plane of the mirror -- a standard approach used to study reaching tasks (Krakauer et al. 1999; Shadmehr and Mussa-Ivaldi 1994; Shadmehr et al. 2010). The targets were presented by an inverted monitor (refresh rate 60 Hz) above the mirror setup which gave the impression that the targets appeared below the mirror while the hand could not be seen. All the experiments were performed using TEMPO/VIDEOSYNC software (Reflecting Computing, St. Louis, MO) that displayed visual stimuli, sampled and stored hand position with other behavioural parameters in real time. The hand positions and joint angles was recorded with a spatial resolution of 0.03 inches using an electromagnetic position and orientation tracking device at 240 Hz (LIBERTY; Polhemus, Colchester, VT).

**Figure 1:**
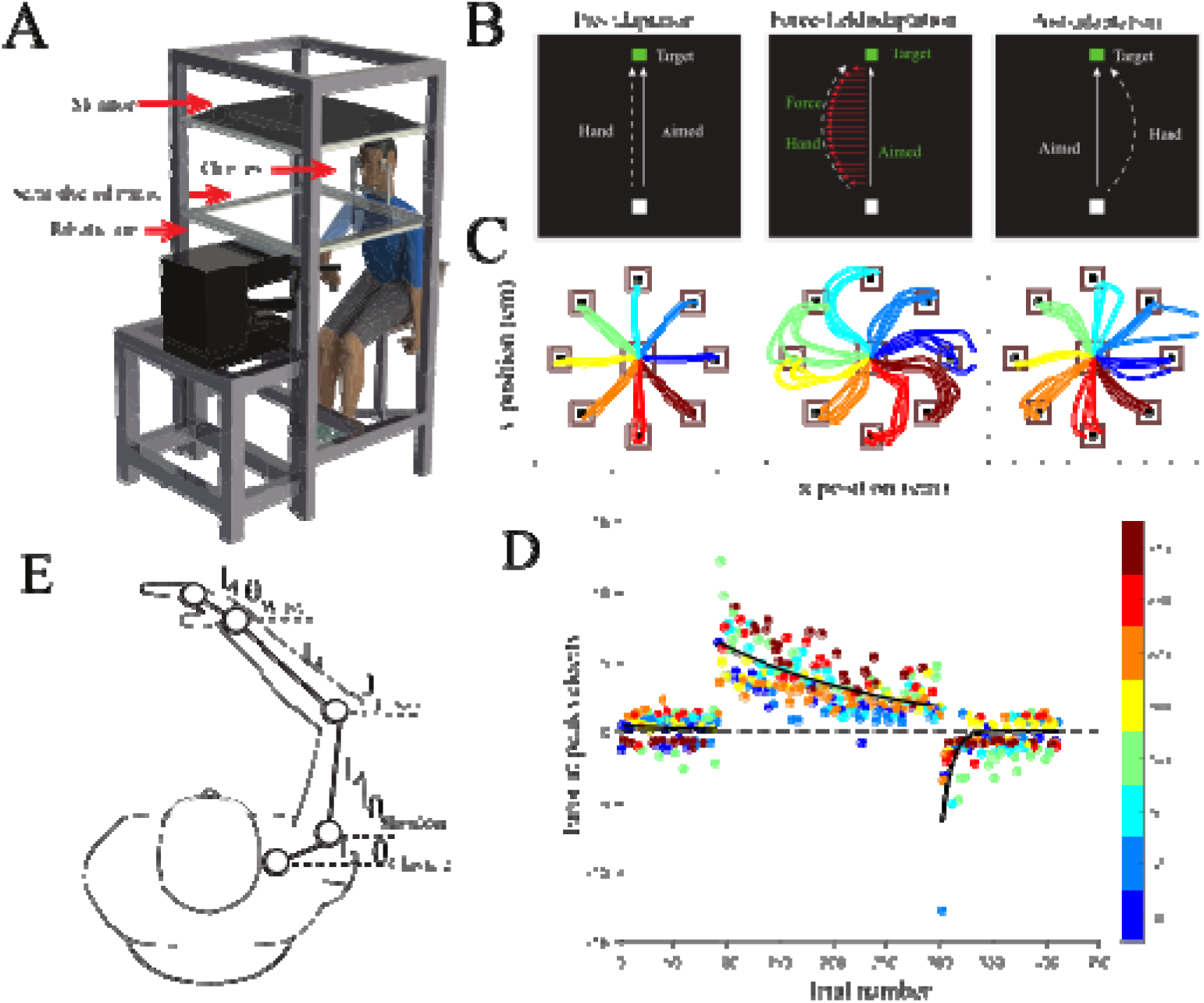
Experiment setup and design: (A) Subjects made point-to-point reaching movements using robotic arm. (B) Subjects made point-to-point reaching movements to visual targets in 1 out of 8 directions 15 cm away from the central start point in each trial. Experiments were divided into three epochs -- a pre-adaptation, an adaptation to an external novel force field and post-adaptation epochs (left, centre and right panels). (C) First five trials of pre-adaptation, force field adaptation, and post-adaptation trials from a subject showing baseline variability, adaptation, and washout effects. (D) Error at peak velocity in pre- adaptation, adaptation, and post-adaptation showing the progression of adaptation for a subject. Errors in each of the eight directions are color-coded. Fitted exponential (black line) significantly accounts for most of the progression of errors across trials during adaptation. (E) Arm model and tracker positions to measure joint rotation angles.

### Experimental paradigm

In all experiment, trials are divided into three phases -- baseline trials, perturbation trials and washout trials. In the baseline portion, the robot arm was free. In the second perturbation trials, the robot applied a force perpendicular to the hand trajectory (discussed in detail below) and in the third phase, the force applied by the robot was switched off. All subjects performed ~30 practice trials before performing the actual experiment. The subjects performed about 400 trials per session with a typical session lasting between 2 to 3 hours. Each trial started with the presentation of a fixation box at the centre of screen. When the robotic end-effector was on the fixation box, then the target box was displayed. The target box was displayed 15 cm away from the fixation box in any one of the 8 directions. The subject moved the robotic end-effector to the target box only after the fixation box disappeared. Till the time fixation box disappeared, subject did not move their hand. Auditory feedback (beep sound) was given when the subject performed correctly.

The top of figure 1 B shows the three phases of the experiment, and the bottom figure 1 C shows the measured first 5 hand trajectories in the three phases for a typical subject. Data similar to figure 1 C was acquired for all trials and across 22 subjects. Figure 1 D shows the error at the peak velocity along the trajectory in the three phases of the experiment for a typical subject. The errors are colour coded -- the blue dots, for example, shows the error when the hand moves along 0 degrees. For each subject, the arm was fitted with electromagnetic trackers which were used to measure joint rotation angles at each instance. The trackers were used to compute the four angles shown in figure 1 E. Additionally, the *(x, y*) location of the end-effector of the robot was also recorded. Detailed analysis of the acquired data for the 22 subjects is presented in the results section.

### Force-field perturbation

During the second phase, the robot applied a lateral force. The lateral force depended on the instantaneous hand velocity as in equation (1)

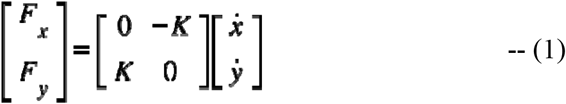

where, are the force components exerted by the robotic arm,, correspond to the velocity of hand and denotes the force perturbation coefficient along the two directions. The force-field disturbs the hand trajectory initially and with trials, the hand trajectory tends to become straighter.

### Visuo-motor perturbation

During the visuomotor perturbation, the cursor movement was rotated according to equation (2),

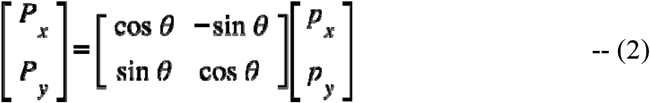

where correspond to the position of the cursor, correspond to the actual position of the hand and (45) denotes the perturbation angle about the center of workspace. This perturbation led to a trajectory error that was gradually compensated over the course of many trials and the hand trajectory straightened. The end-point curser describing the movement was visible throughout the movement in all experiments.

### Kinematic model of arm

A forward kinematics model of the human arm is assumed to have four joint rotations -- clavicle protraction–retraction, shoulder horizontal abduction–adduction, elbow flexion–extension and wrist medial–lateral -- denoted by respectively (see figure 1 E). The Cartesian location of the hand can be written as

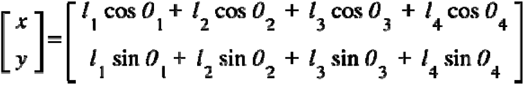

where the four joint rotations are as described above. It may be noted that the angles, i = 1, 2, 3, 4 are absolute angles and hence the forward kinematic equations above are different from typical serial robot kinematic equations with relative angles. The link lengths, i=1, 2, 3, 4 are computed from the data from the sensors placed in the arm and vary a little with different subjects. To ensure that the values are valid, the (*x, y*) obtained from above equation is compared with the (*x, y*) values of the robotic arm handle and the values are determined such that the difference was less than 1.0%

Based on the forward kinematic model, the Jacobian matrix at any joint configuration, can be obtained as

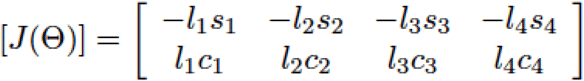

where is the vector and *c*_i_ and *s*_i_ denote cosine and sine of angle The Cartesian error is related to the joint error as

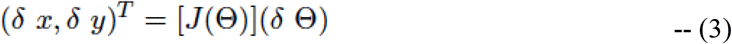

It can be seen that the Jacobian matrix depends on the 4 joint variables and hence will vary at different points in the workspace. For a unit, the maximum and minimum are the maximum and minimum singular values of and they occur along the mutually orthogonal singular directions, and the unit circle maps to an ellipse in the Cartesian space. If the Jacobian matrix is independent of, scaling can be used to make the minimum and maximum singular values equal and thus the error distribution can be a circle (Raibert and Craig 1981).

### Quantifying redundancy in the arm

In a serial robot, the motion of the end-effector |(∂*x*, ∂*y*)^*T*^| is not affected by joint motion ∂ Θ, then the robot is said to be redundant. In the arm model, the number of joint variables is 4 while the Cartesian motion is of dimension 2. This implies that there exists some Θ which do not affect the Cartesian motion and there are redundant degrees of freedom in the arm. The redundancy in the arm model is quantified by *N(J)* obtained from the null space of Jacobian matrix, [*j*(Θ)], as follows:

The variability in the joint variable, Θ, is obtained for the baseline trials for each of the 8 different directions at the maximum Cartesian velocity or at the maximum error along the trajectory. The mean joint configuration across trials, Θ, was computed also at the maximum Cartesian velocity. The joint configuration for the *k*^th^*trial* Θ_k_ was subtracted from the mean to obtain the deviation, ΔΘ_k_ as

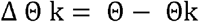

To compute the null space of the Jacobian matrix, it is assumed that the mean Θ results in the nominal hand trajectory and a part of deviation ΔΘ_k_ are in the null space of [*J*(Θ)]. The vectors in the null space of the Jacobian matrix, *ξ_i_*, are obtained from

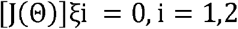

and the component of ΔΘ_k_ lying in the null space spanned by {*ξ*_1_, *ξ*_2_} are the inner products < ΔΘ_k_, *ξ_i_* >, *i* = 1,2. The sum of the two null space components for the k^th^ trial is computed as

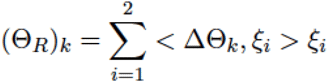

and *N*(*J*) is defined as

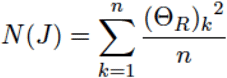

where *n* is the number of trials.

### Quantifying learning

One of the simplest models of learning is that of a first-order process where the error is reduced exponentially. The variation of peak error *e(t)* as a function of time *t*, can be written as

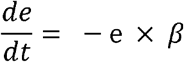

where β is a parameter that describes the rate of change of error and is independent of the current error. The evolution of error from the above can be written as.

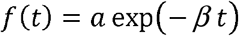

where α is the initial error and β is the learning rate.

In our case, the error is the maximum deviation (perpendicular distance) of the hand from the straight line connecting the fixation box and the target box in each of the 8 directions. Additionally, instead of a continuous function of time, the error is for each trial, and we have

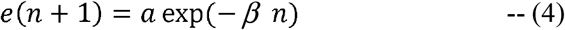

where *n* denotes the trial number.

### Markov Chain Monte Carlo Method

During the experiments, the subjects performed approximately 200 trials in the 8 directions. The error in the 8 directions is sorted into columns and on an average, there are 25 data points for each direction -- they vary between 17 and 39 dues to the random nature of the chosen target directions. A typical table of error in centimetres in each trial along each of the 8 directions is shown in Supplemental Table 1 for the subject whose data is shown in figure 1 D.

To obtain α and β for each of the directions, we used a Markov Chain Monte Carlo (MCMC) approach (Suess and Trumbo 2010). The main motivation for using a MCMC method is that it is known to be more robust in comparison to other nonlinear curve fitting schemes. It also provides the distribution of the parameters for a chosen prior distribution -- chosen as same uniform distribution for all directions and subjects in this work-- giving a much better insight and confidence to the values obtained for α and β. The values of α and β obtained using MCMC for the subject above along direction 90 degree (column 4 in Table above) is shown in Supplemental Table 2. The values of α and β obtained using the well-known Levenberg-Marquardt (LM) nonlinear curve fitting scheme (Nocedal and Wright 2006) is also shown below for comparison, and the numbers obtained from both the methods are in good agreement. It may be mentioned that the R language was used to implement the MCMC and Levenberg-Marquardt algorithms.

The Supplemental Table 2 shows the values of α and β obtained using the LM and the MCMC approaches which indicate that mean values of α and β obtained are consistent. The main advantage of MCMC over LM is that we also obtain the distribution of α and β which gives a much better confidence to the obtained numbers. We have used MCMC for all 22 subjects and for all 8 directions.

## Results

We trained 12 subjects to learn point-to-point reaching movements using their dominant hand, along 8 directions, in a force-field which was set using the force-field perturbation equation (1). In this experiment the perturbation was proportional to the velocity of the hand but perpendicular to the hand movement direction. We used an experimental setup shown in figure 1 A. The data for the three phases – a pre-adaptation baseline period, followed by a force field perturbation (dynamic perturbation) and finally a post-adaptation phase when the perturbation was removed – figure 1 B for a representtaive subject. The data consists of the location of the end point and joint angles while subjects reached/pointed to the target during the three phases.

Figure 1 C shows the trajectories for the first five trials in the three phases for the representative subject. The peak error was calculated as the perpendicular distance of the hand trajectory at peak velocity from the straight line joining the central fixation box to the target location. Overall, the pattern of trajectories are consistent with previous work showing that while typical movements follow a straight trajectory in the baseline condition, they show strong curved trajectories in the presence of a force-field. The curved trajectories gradually become straighter with practice over the course of about two hundred trials with the error decreasing as shown in figure 1 D. In addition, as a consequence of motor learning, subjects showed a washout effect (post adaptation) where errors in trajectory inverts in direction when the learned force field is turned off in the post adaptation period. This washout error converges to baseline levels.

We assume that learning is a first-order process where the error is reduced exponentially, and the variation of peak error is as shown in equation (4). To compute the learning in force--field perturbation trials, the errors were fitted with an exponential fit using the Levenberg-Marquardt algorithm, and for the representative subject β = 0.0064 ± 0.0006 Std. error, *t* = 11.51 and *Pr* (> | *t* |) < 2*e* - 16 and *a* = 6.41 ± 0.292 cm Std. error, *t* = 21.92 and *Pr* (> |*t* |) < 2e - 16(see Figure 1D).

To obtain a more geometric view, we plot the mean error and variation in the baseline, first 5 trials and last 5 trials under force-filed perturbation in the 8 directions. An ellipse is fitted (motivated by the velocity distribution at a point seen in a robot -- (see text after equation (3)) with the mean error in the 8 directions for each of the three data sets. The mean error is denoted by a “circle” and the “line” through the “circle” denote the variation with trials. Figure 2 A shows the error distribution along 8 directions for the representative subject data shown in figure 1. The size of the error ellipse decreases from the first 5 to the last 5 trials -- the errors are large when the lateral force is applied and as the trials progress, the error decreases, indicating that the subject learns to adapt to the external lateral force. It can be observed that the error ellipse in the baseline (when no external force is applied) is the smallest.

**Figure 2:**
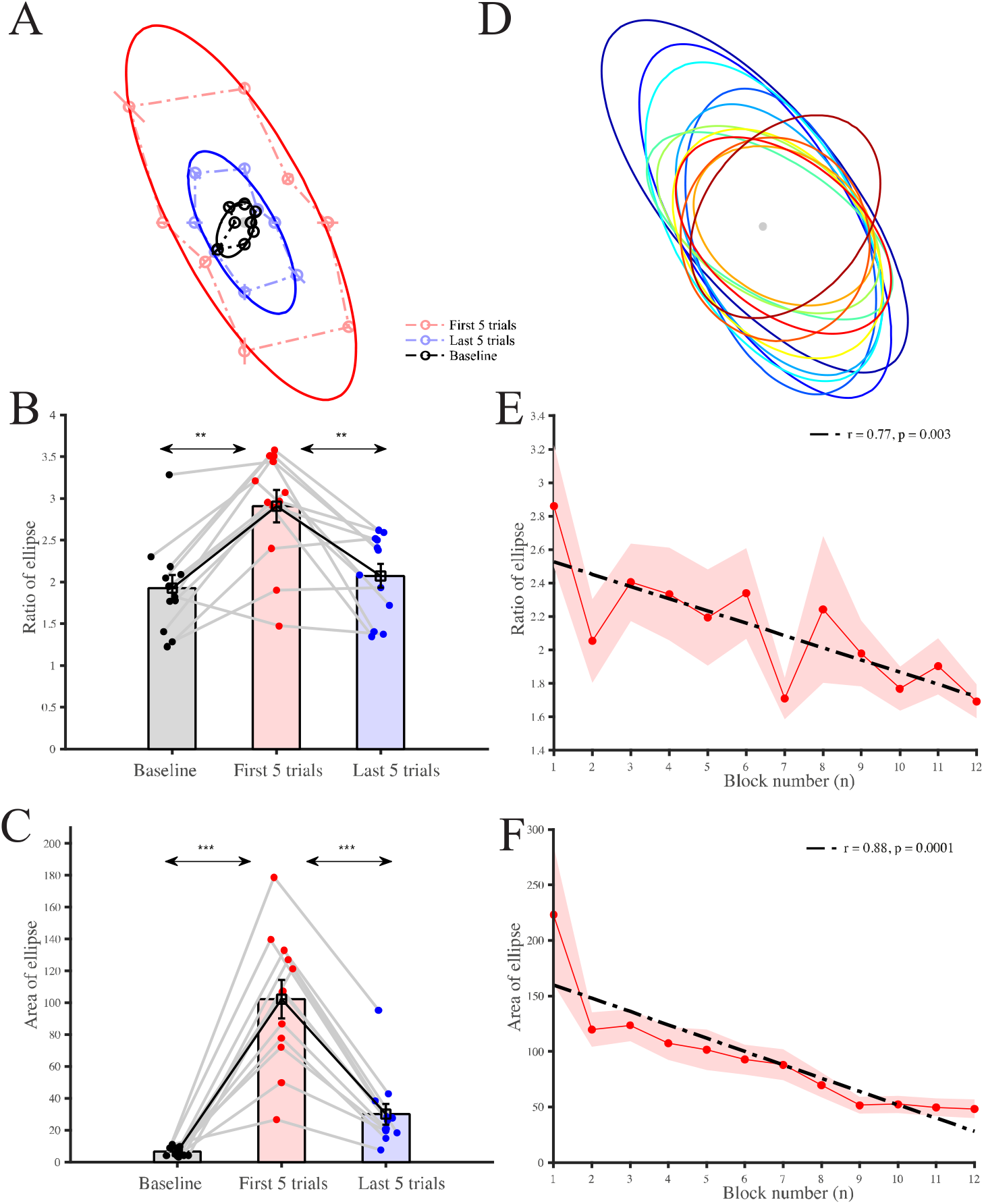
Homogenization of workspace in the dominant arm: (A) Representative subject directional error ellipse for the baseline (black), starting of perturbation (red) and end of perturbation (blue). (B) Comparison of directional error ellipse eccentricity in baseline (black), starting of perturbation (red) and end of perturbation (blue) reveal higher eccentricity in the starting of perturbation. (C) Comparison of directional error ellipse area in baseline (black), starting of perturbation (red) and end of perturbation (blue) reveal higher area in the starting of perturbation. (D) Representative subject directional error ellipse progression towards a more circular ellipse. The ratio gradually decreases with practice over the course of about two hundred trials (E) Regression across subjects showing the progression of change in ellipse eccentricity indicative of homogenization of error distribution. (F) Regression across subjects showing the progression of change in ellipse area indicative of learning.

Figure 2 B and C shows the bar plot of the ratio of the major to the minor axis and the area of the ellipse for all the subjects. The ratio is large in the first 5 trials and the ratio in the last 5 trials approaches the value in the baseline. The mean ratio in baseline period (mean = 1.93 ± 0.55) was significantly less than the mean ratio at the starting of perturbation (mean = 2.91 ± 0.67) (figure 2 B; *p* = 1.44e-4, *t* (11) = 5.67). There was also significant difference in the mean ratio between the starting of perturbation and end of perturbation (figure 2 B; mean = 2.07 ± 0.50; *p* = 0.002, *t* (11) = 4.16). Figure 2 D shows the evolution of the error ellipse with trails for subject.

We also computed the area of the ellipse, which is a proxy for overall learning rate, across directions. The area in the baseline period (mean = 6.44 ± 2.68) was significantly less than the mean area at the starting of perturbation (mean = 102.23 ± 42.02) (figure 2 C; *p* = 8.25e-06, *t* (11) = 7.80). There was good difference in the mean ellipse area between the starting of perturbation and the end of perturbation (figure 2 C; mean = 30.02 ± 22.68; *p* = 1.82e-06, *t* (11) = 9.13). However, the area of the ellipse is larger in the last 5 trials as compared to the baseline in all the 12 subjects. The area of the ellipse gradually decreases with practice over the course of about two hundred trials as shown in figure 2 F.

Overall, the main findings are that the maximum error due to the external force-field is very large when it is applied and due to learning the error decreases with trials. This can be seen from the straightening of the curved trajectories as shown in figure 1 D for a subject and quantitatively in the plot of area of the ellipse for 12 subjects shown in figure 2 D. Secondly, as shown in figure 2 E, the ratio of the major to minor axis of the ellipse decreases with trials and the error distribution tends to become circular -- it is not a perfect circle which would imply a ratio of 1.0. The learning results shown by the 12 subjects is not only in terms of reduction of error magnitude but also making the error distribution more uniform -- this is termed as homogenization of the task space error distribution.

### Active learning of a force-field perturbation

Differences in the ratio of the major to minor axis may reflect a difference in the intrinsic biomechanics of the human arm. In contrast, differences in the ratio may also reflect the effect of neural control that assists in homogenizing the workspace of the human arm. To assess this, we tested and compared the ratio of the major to minor axis of the ellipse across direction in dominant and non-dominant hand in 10 subjects, thereby normalizing any differences in the biomechanics.

Similar to the dominant hand, the figure 3 A shows the measured maximum peak error in the baseline where no perturbation is applied, in the first 5 and last 5 trials with force-field perturbation in each of the 8 directions with the non-dominant hand for a typical subject. Again, the mean error is denoted by a “circle” and the “line” through the “circle” denote the variation with trials. We fit ellipses through the mean error along the 8 directions. This is shown in figure 3 A for the baseline, the first 5 and the last 5 trials. It can be seen that as trials progress the ratio of major to minor axis in non-dominant hand do not decrease as seen in the dominant hand (see figure 2 E). The mean ratio in the baseline period (mean = 1.63 ± 0.41) was significantly less than the mean ratio at the starting of perturbation (mean = 2.15 ± 0.50) (see figure 3 C; *p* = 0.03, *t* (9) = 2.53). There was no difference in the mean ratio between the starting of perturbation and end of perturbation (see figure 3 C; mean = 2.16 ± 0.81; *p* = 0.97, *t* (9) = 0.03).

**Figure 3:**
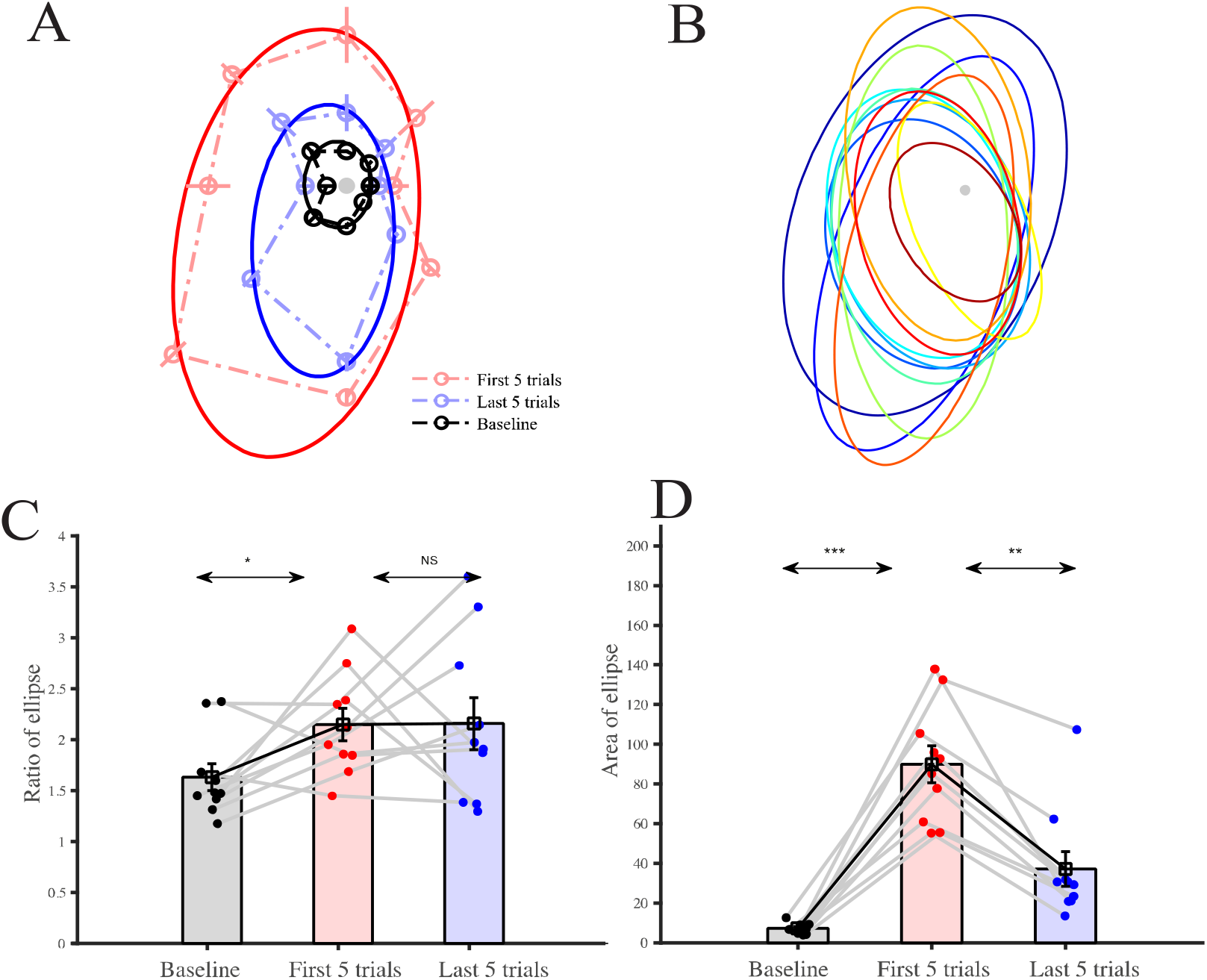
Homogenization of workspace in non-dominant arm: (A) Representative subject directional error ellipse for the baseline (black), starting of perturbation (red) and end of perturbation (blue). (B) Representative subject directional error ellipse progression. (C) Comparison of directional error ellipse eccentricity in baseline (black), starting of perturbation (red) and end of perturbation (blue) reveal higher eccentricity in the starting of perturbation. (D) Comparison of directional error ellipse area in baseline (black), starting of perturbation (red) and end of perturbation (blue) reveal higher area in the starting of perturbation.

The mean ellipse area in baseline period (mean = 7.19 ± 2.66) was significantly less than the mean ellipse area at the starting of perturbation (mean = 89.94 ± 29.43) (see figure 3 D; *p* = 5.83e-06, *t* (9) = 9.43). There was significant difference in the mean direction ellipse area between the starting of perturbation and end of perturbation (see figure 3 D; mean = 37.07 ± 27.90; *p* = 1.39e-04, *t* (9) = 6.31). The ellipse area gradually become smaller with practice over the course of about two hundred trials which implies some learning is taking place.

Taken together these findings indicates that the difference in ratio of the major to the minor axis between the dominant and non-dominant hand or the capability of homogenization of the error distribution maybe a consequence of the different mechanisms in the dominant hand and non-dominant hand -- most likely due to active neural control.

### Maximum error, learning rate and redundancy along directions

As mentioned earlier, the learning rate β and the maximum error α see equation (4) for all the 200 trials (along all directions taken together) for a particular subject was 6.40 and 0.006. Similar results were obtained for all the 12 subjects. To investigate the variation of α and β along each direction, the force-field perturbation trail data are sorted along the 8 directions. This is shown for a typical representative subject is shown in Supplemental Table 1-- the variation from 25 trials along each direction is due to the random nature of the target presented. To obtain α and β in each of the 8 directions, we used the robust MCMC algorithm. The mean α and β values across 12 subjects is shown in figure 4 A and C and obtained values in Supplemental Tables 3 and 4 (in Supplemental Table 3 and 4, the values of α and β for subject 9 is not considered in the analysis since the results obtained from MCMC and LM differ significantly.), respectively. Figure 4 of B and D shown the bar plots of α and β across subjects. There is statistical difference in α value in each of the directions (F (7,79) = 5.54, *p* = 3.24e-5, see figure 4 C) indicative that some directions have higher initial errors. Furthermore, there is a clear statistical difference in learning rate along 90° and 225° (F (7,78) = 3.11, *p* = 0.006, see figure 4 D). These are also very close to the major and minor axis of the ellipse of error distribution.

**Figure 4:**
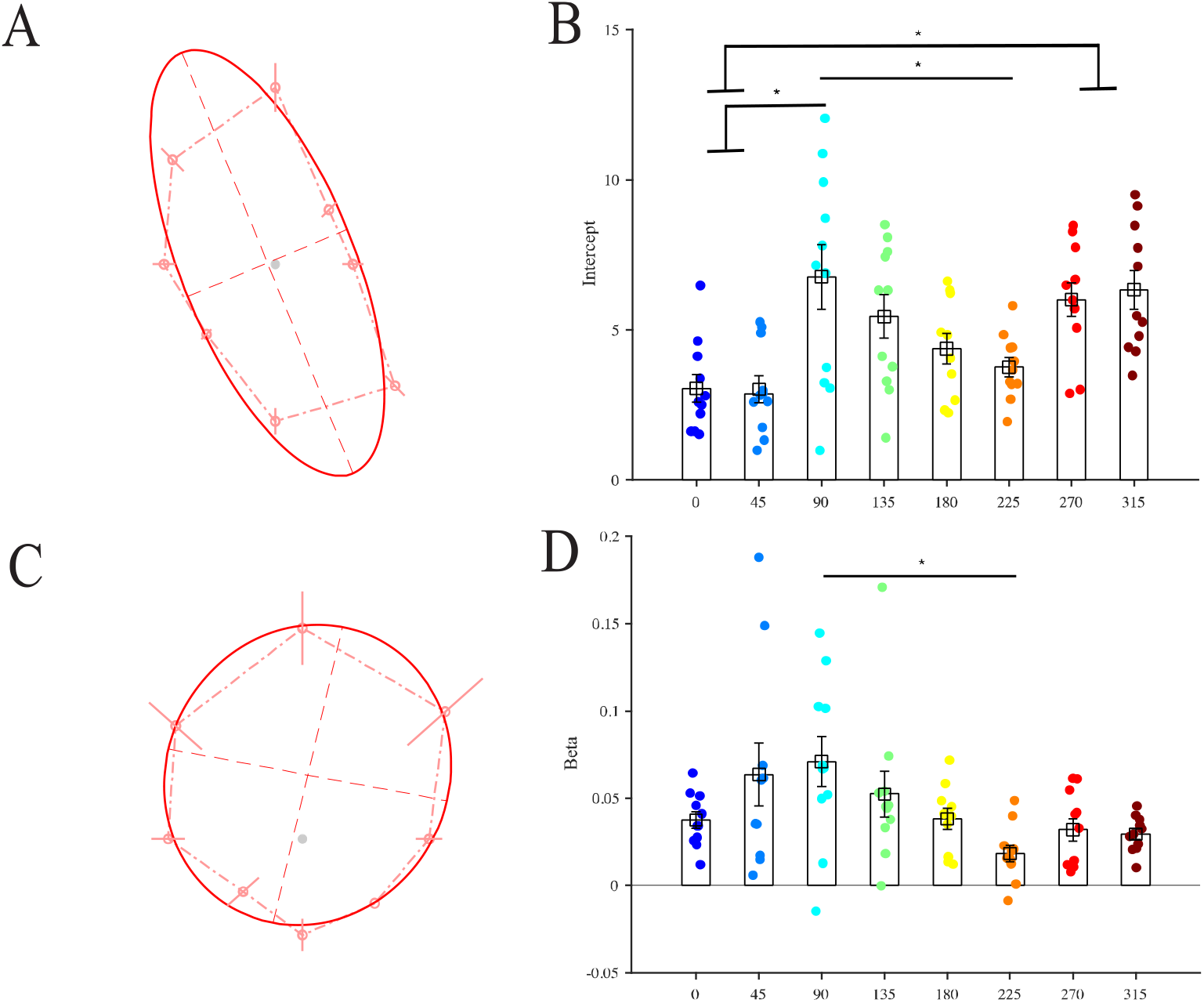
Learning is greater for more difficult directions leading to homogenization. Difficult direction in adaptation: (A) Intercept mean ± se across subjects showing the ratio of the major to minor axis of the ellipse indicative of some direction has higher initial errors. (B) Comparison of intercept in different directional. (C) learning rate mean ± se across subjects showing the learning rate in different directions. (D) Comparison of learning rate in different directions.

As mentioned earlier, the human hand is known to be redundant. In the model shown in section material, a four degree-of-freedom model was assumed for the human hand performing planar point-to-point reaching motions and hence, the human hand is assumed to possess two redundant degrees of freedom. In order to test whether redundancy could aid in homogenization of error distribution due to a force-field perturbation, we computed the null space of the Jacobian *N(J)*, representing the use of the redundancy in the four degree-of-freedom arm model. Figure 5 shows a bar plot of *N(J)* for the dominant and non-dominant hand for all subjects in the baseline period. Consistent with our hypothesis, *N(J)* was lesser along the minor axis (mean = 0.06 ± 0.05) compared to the major axis (mean = 0.11 ± 0.07, *p* = 0.006, *t* (9) = 3.49,) for the dominant hand. For the non-dominant hand, there was no difference in *N(J)* values along the major and minor axis.

**Figure 5:**
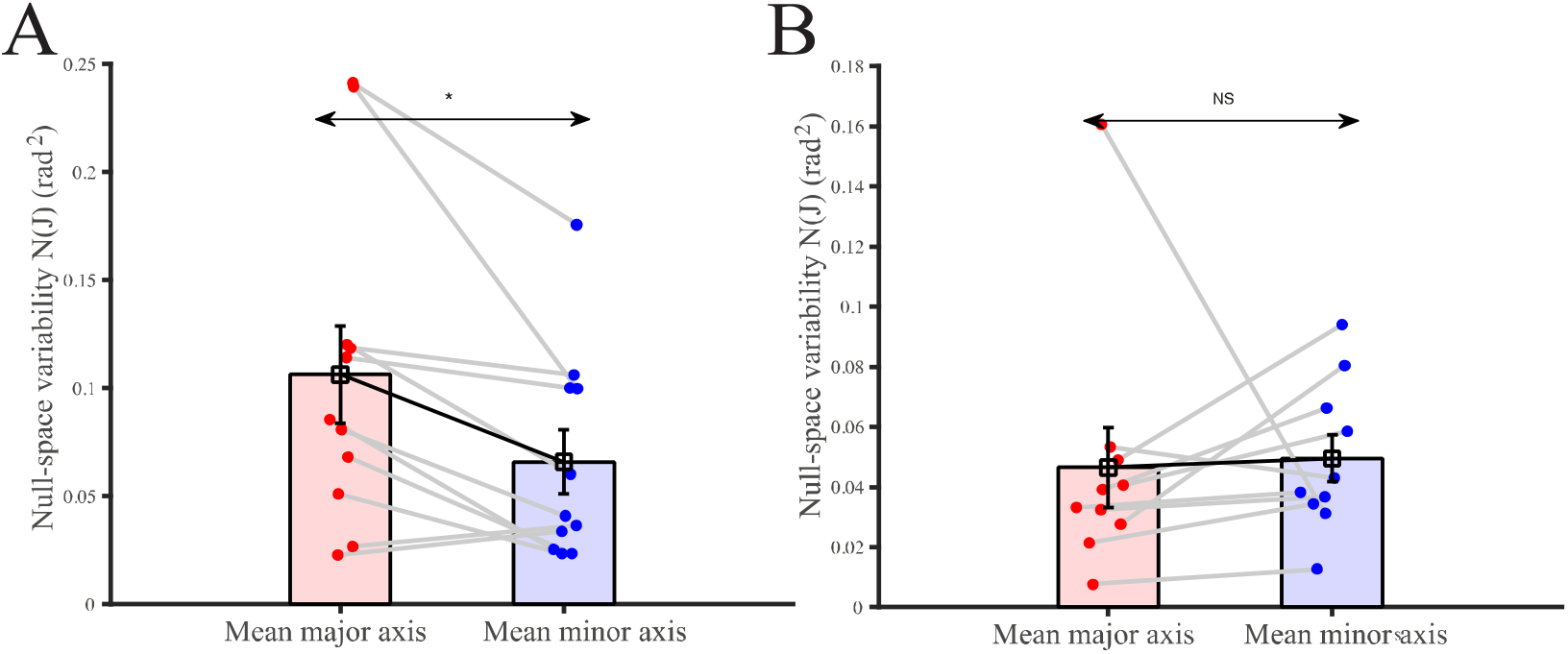
Null-space variability in the major axis and minor axis in direction ellipse. (A) Comparison of dominant hand null-space variability along major axis(red), and minor axis (blue). (B) Comparison of non-dominant hand null-space variability along major axis(red), and minor axis (blue).

We have earlier observed that during learning homogenization of the error takes place, i.e., the ratio of the major to minor axis of the direction error ellipse decreases with trials. Taken together we can suggest that redundancy may also play a role in making error distribution more uniform, i.e., use of redundancy leads to homogenization of error distribution in point-to-point reaching tasks.

### Learning of a visuo-motor perturbation

To test whether homogenization is a property related specifically to learning of the newly imposed biomechanics or is a more general feature of motor learning we next examined 10 subjects while they learnt point-to-point reaching movements with the visuo-motor perturbation, equation (2), along 8 directions, where in each case, the cursor was rotated by 45° from the hand trajectory. Similar to the dominant hand, figure 6 A shows the measured maximum peak error in the baseline where no perturbation is applied, in the first 5 and last 5 trials with visuo-motor perturbation in each of the 8 directions with the dominant hand for a typical subject. Again, the mean error is denoted by a “circle” and the “line” through the “circle” denote the variation with trials. We fit ellipses through the mean error along the 8 directions. This is shown in figure 6 A for the baseline, the first 5 and the last 5 trials. Unlike in the dynamics condition, as trials progress the ratio of major to minor axis in visuo-motor perturbation and did decrease for the dominant hand (see figure 2 E). The mean ratio in the baseline period (mean = 2.41 ± 0.71) was different than mean ratio at the starting of perturbation (mean = 1.59 ± 0.29) (see figure 6 C; *p* = 0.018, *t* (9) = 2.88). There was also no difference in the mean ratio between the starting of perturbation and end of perturbation (see figure 6 C; mean = 1.89 ± 0.55; *p* = 0.18, *t* (9) = 1.46).

**Figure 6:**
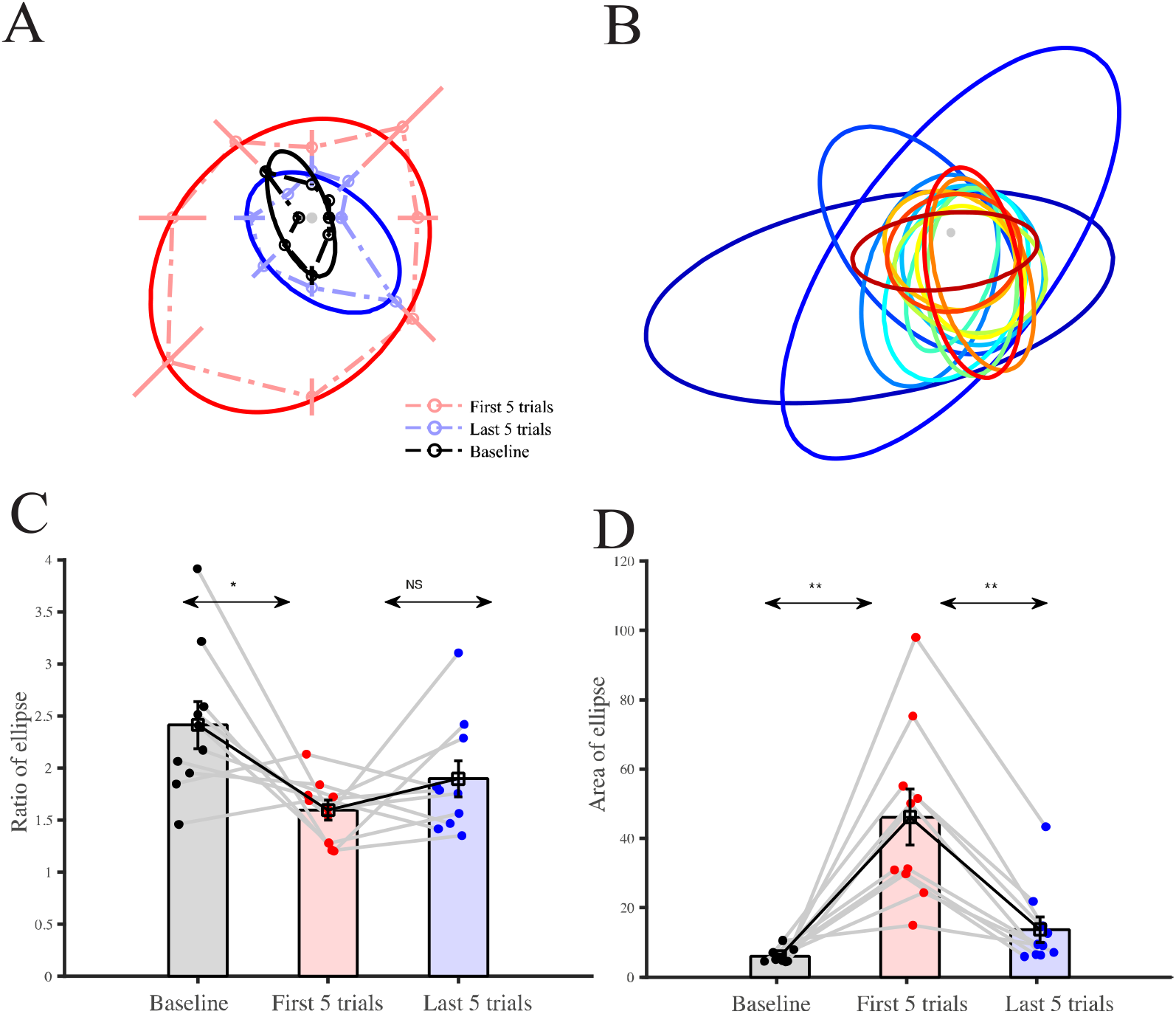
Learning in the absence of homogenization of workspace during visuomotor adaptation: (A) Representative subject directional error ellipse for the baseline (black), starting of perturbation (red) and end of perturbation (blue). (B) Representative subject directional error ellipse progression. (C) Comparison of directional error ellipse ratio in baseline (black), starting of perturbation (red) and end of perturbation (blue) reveal a higher ratio in the starting of perturbation. (D) Comparison of directional error ellipse area in baseline (black), starting of perturbation (red) and end of perturbation (blue) reveal a larger area in the starting of perturbation indicating learning

The mean ellipse area in baseline period (mean = 6.06 ±1.95) was significantly less than the mean ellipse area at the starting of perturbation (mean = 46.15 ± 25.44) (see figure 6 D; *p* = 0.001, *t* (9) = 4.76). There was, however, a significant difference in the mean direction ellipse area between the starting of perturbation and end of perturbation (see figure 6 D; mean = 13.73 ± 11.52; *p* = 2.06e-4, *t* (9) = 5.98). The ellipse area gradually become smaller with practice over the course of about two hundred trials as shown in figure 6 D which implies learning. occurs in the absence of homogenization of the error distribution in the visuo-motor perturbation condition.

## Discussion

In this study, we have presented variation in task space error along directions in point-to-point reaching tasks. We demonstrated that the motor learning is homogenizing the workspace of the human arm possibly to increase the efficiency and accuracy. In addition, our results also showed a significantly larger use of redundancy along the directions with more error (or major axis) versus the direction with less errors (or the minor axis) of error ellipse. The data revealed novel direction specificity of motor learning not reported earlier to the best of our knowledge.

What is more significant is the observation that this anisotropy in errors distribution is reduced with trials and the error distribution becomes more circular. This work also indicates that the redundancy in the arm is used to homogenize the error distribution. Although not known where in the central nervous system the resolution of redundancy is performed and how, this works provides a possible reason on how the redundancy in the arm is used for reaching tasks.

### Homogenization and Linearization of Control

In control theory, it is well known that it is significantly easier to design controllers for a linear system to achieve a desired accuracy. More specifically, for a robot to follow a desired path with desired accuracy, extensive effort has gone into designing controllers starting from the simplest proportional, integral and derivative (PID) to sophisticated model-based control (Craig 2004). A robot can be considered to be linear up to first order if the Jacobian matrix relating the end-point velocity to the joint rates is constant and for such a Jacobian matrix, the error distribution is circular or isotropic. From control theory, it has been argued that such a “linear” robot would be easier to control and would achieve better accuracy -- such robots containing only sliding joints and with no rotary joints was attempted in the earlier days of robotics but was given up due to the issues in friction at the sliding joints. This work indicates that the learning leads to approximate homogenization of the error distribution, i.e., makes the arm more ``linear’’ even though it contains rotary joints which makes the Jacobian matrix a function of the location and inherently nonlinear. A possible consequence of homogenization of the error distribution would be that the overhead on the central nervous system in terms of control during reaching tasks is reduced.

### Homogenization and Biomechanics

Our results indicate that some directions appear to be easier to learn and some are more difficult. In Howard et al. 2013 (see also Singh et al. 2016), error plots in 8 directions indicate some dependence on direction but this is not brought out clearly. In this work, we can clearly see that the errors are significantly larger in direction of the major axis and significantly smaller in the direction of the minor axis -- the mean (across subjects) major axis is observed to be around 111.68°, and the minor axis is around 201.68°. It is known from biomechanics of human arm (or dynamics of serial robots) that the inertia of the arm (as seen from shoulder) and the effort required is largest when the arm is moving in the direction of the major axis and likewise the inertia seen, and the effort required is smaller when the arm is moving along the minor axis and a mechanistic view of more/less error along direction of more/less effort is consistent with this observation.

The task space positioning error for a robot, in the presence of external disturbances and noise, is also related to the stiffness (or impedance) of the robotic arm. The impedance of the robot arm is typically determined by the control scheme (Hogan 1984; Raibert and Craig 1981) and controller gains, and the larger the stiffness the less is the positioning error. In reaching tasks by human arms, position control involves increase in limb impedance (Franklin et al. 2007; Wong et al. 2009). In (Wong et al. 2009), the authors claim that the limb stiffness is modulated to achieve accuracy requirement in the absence of destabilizing force, and they observe a modest increase in limb stiffness perpendicular to the direction of motion when more accuracy is required while moving along a narrow track. In (Franklin et al. 2007) the authors state that the end-point stiffness, primarily due to the co-activation of bi-articular muscles, approximately aligns with the direction of instability in the environment with the stiffness ellipse rotating to align with the direction of instability. In this work, we proceed further and show that learning results in homogenization, possibly through changes in the stiffness properties as well as changes in the internal model of the arm.

### Homogenization is controlled by the CNS

A second major observation in this study is that bio-mechanically similar dominant and non-dominant arms showed different homogenizing behaviours. Both the dominant and the non-dominant hand shows directional dependency of error and both show that the size of the error ellipse decreases with trials. This indicates that learning is present in both arms. However, the homogenization of error distribution is not seen in the non-dominant hand and this indicates that homogenizing is being actively done by the central nervous system -- the homogenizing of the workspace seen in the dominant hand provides a natural explanation of why learning might be more potent in the dominant hand and reaffirms the belief that homogenizing not only reflects the bio-mechanical characteristics of the arm but may reflect active control from brain or the central nervous system. In a related set of experiments with visuo-motor rotation, all directions show the same error distribution (see figure 4). This is consistent with our study as learning a visuo-motor rotation involves learning of an internal kinematic model and unlikely requires the homogenization since the inhomogeneities are relate to dynamics.

### Homogenization by feedback control internal models

As mentioned previously, movements associated with larger inertia will have greater muscle loads and as a consequence of signal dependent noise will generated larger errors. In this context, it is interesting that the distribution of errors in the baseline condition is low and homogenous despite inhomogeneous biomechanics. Such uniformity is likely to be a consequence of kinematic feedback controllers that ensure of uniform endpoint control, as well as internal models that ensure homogeneity even during the early feedforward driven aspect of the trajectory. This suggests that in the baseline the brain (CNS) uses a well learnt sensorimotor mappings that get transiently inactivated, exposing the underlying inhomogeneous biomechanics. Although, we do not propose a mechanism on how feedback gains and internal models are relearnt following the perturbation, our results suggest that a form of learning that is sensitive not just to the magnitude of the errors as a consequence first order learning, but of the learning rates themselves that are sensitive to the direction of movement that helps homogenise errors. We suggest that such directional specific learning linearizes responses and facilitates generalization beyond the region of training One important issue is direction generalization function – this refers to the movement direction that displays the maximum degree of adaptation after learning with the degree of adaptation falling off with adjacent movements. Donchin et al. (2003) argue from experiments conducted during reaching tasks, subjected to a force-field disturbance, that the generalization is bi-modal, perhaps reflecting basis elements that encode direction bi-modally. This work also does not discuss the direction dependence of positioning error in reaching tasks. In this work, our results indicate that the generalization is not bi-modal -- the distribution of error is best approximated by an ellipse with some direction showing large error and some direction showing less error across subjects.

### Homogenization, redundancy, and learning

In previous work we showed through first and second order correlations of null space variability--a proxy for joint redundancy--a possible role for joint redundancy in motor learning of dynamics and kinematics. Here, we extended this correlation to study the directional dependency of redundancy and its possible impact on motor learning. In congruence and extension with our previous study, we observed that the redundancy exhibited a directional axis that aligned with the learning axis, which could also explain the observed homogenization. Furthermore, this redundancy axis was only observed for the dominant arm and not seen in the non-dominant arm which suggests that this alignment maybe causal in nature. In contrast to the learning of dynamics, kinematic perturbations did not produce inhomogeneities in errors for redundancy to exploit. Taken together, we speculate that a greater redundancy may allow better learning by increasing available options, contributing to homogenising errors across directions.

## Conclusions

This paper deals with motor learning in point-to-point reaching tasks performed by a human arm. Experiments conducted with adult subjects show that maximum error along the trajectory, when the arm is perturbed by a lateral force, decreases with practice and approaches the situation when no lateral force is applied. In this paper, we investigate the error in the 8 different directions in which the reaching task was carried out. The error distribution along the 8 directions is fitted with an ellipse and it is observed that, across subjects, a direction between 90 and 135 degrees has the largest error while a direction between 180 and 225 degrees shows the least error. As the trial progresses and learning takes place, the magnitude of the error along all directions reduce to a value close to the baseline trials where no lateral force is applied. Moreover, it is observed that the eccentricity of the error ellipse reduces, and the error distribution becomes more circular. These two observations indicate that the learning leads to homogenization of the trajectory error. Similar experiments done with the non-dominant and bio-mechanically similar arm, show that while there is learning (error magnitude decreasing with trials), the eccentricity of the error ellipse does not reduce. This leads us to the conclusion that the homogenization is a consequence of active neural control. Furthermore, analysis suggest that the redundancy in the arm is used to make the error in different directions more uniform and is thus a possible use of the redundancy in all human arms.

A typical anthropomorphic robot is known to have a nonuniform (ellipse) error distribution at different locations in its workspace and in a redundant robot, the redundancy can be used to make the error distribution uniform (circular) in all directions at a location. Uniform error distribution is also seen in a linear system which is known to be more easily controllable. The homogenization result and its analogy with a mechanical robot suggests that the redundancy is used to make the human arm more linear which in turn make it easier for the central nervous system to control. More work is required to obtain a better understanding where the redundancy in human arm is processed and how the redundancy in the actuation system, namely muscles, are resolved.

## Acknowledgment

We thank Dr. Sumitash Jana for the initial setup of the experiment.

## Author Contributions

P.S., A.G. and A.M. designed the experiments, P.S. performed the experiments, P.S. and O.G. analysed the data and P.S., A.G., and A.M. wrote the paper.

## Data Availability

Data available on request from the authors.

