## Supplementary figures and images for "Motor learning in reaching tasks leads to homogenization of task space error distribution"

### Supplemental Table

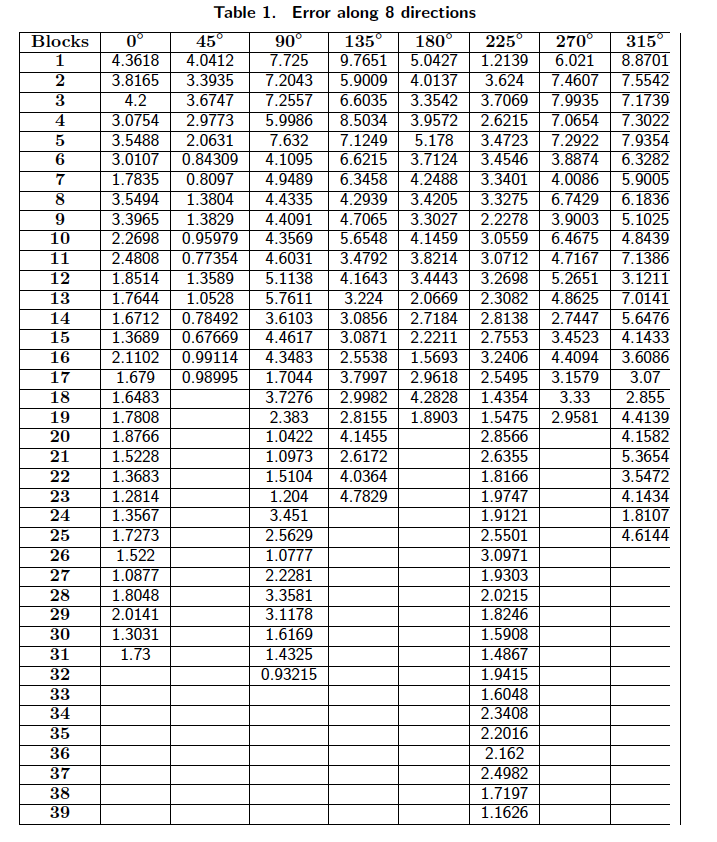

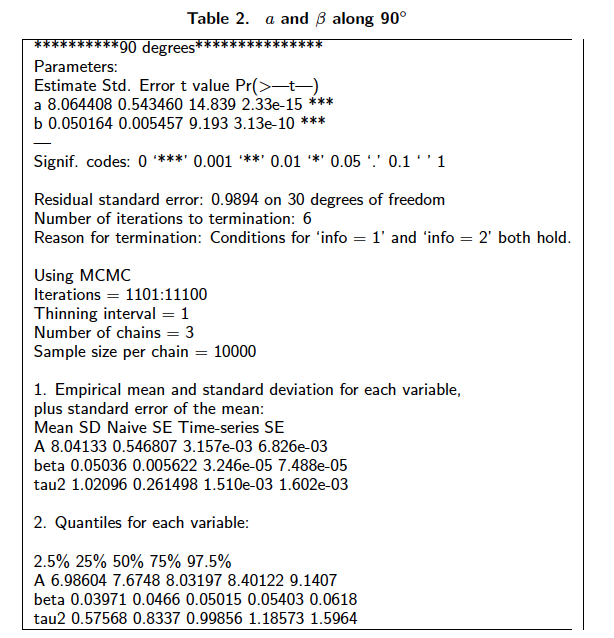

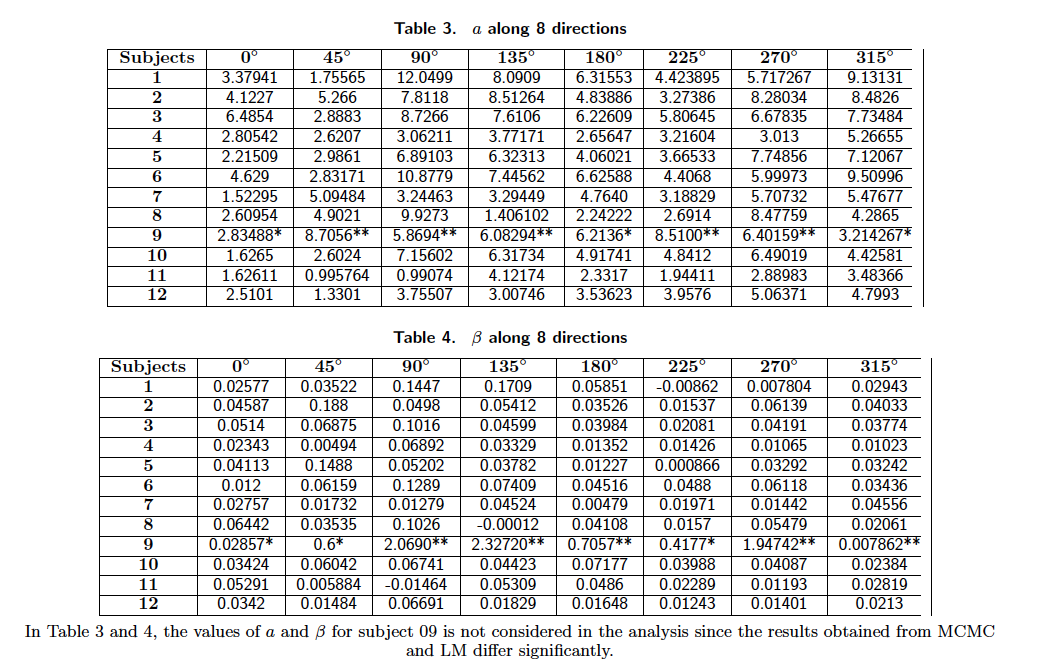
